# In vitro fertilisation and vitrification disrupt embryo mitochondrial function and redox balance that persists into adulthood in mice

**DOI:** 10.64898/2026.08.14.744765

**Authors:** Yutong Chen, Henrietha N Chukwuefe, Min Zi, Gina J Galli

**Author notes:** Corresponding author: Dr Gina J Galli Division of Cardiovascular Sciences School of Medical Sciences, Faculty of Biology, Medicine and Health The University of Manchester Core Technology facility, 46 Grafton Street, Manchester, M13 9NT, UK.

## Abstract

**Background and aims:** Assisted reproductive technologies (ART), including in vitro fertilisation (IVF), account for over 10 million births worldwide. ART-conceived young offspring show altered cardiovascular phenotypes, including cardiac remodelling and raised blood pressure, but the mechanisms remain unclear. Mitochondrial disturbance during preimplantation development may link early ART exposure to later cardiac dysfunction. However, to our knowledge, no one has assessed mitochondrial function in adult offspring from IVF pregnancies. In this study, investigated the effects of IVF and embryo vitrification on blastocyst mitochondrial redox balance and metabolism, and determined whether these effects persisted into the adult heart.

**Methods and Results:** IGS-CD1 mouse blastocysts from naturally mated donors or IVF were transferred fresh or after vitrification–warming. IVF reduced blastocyst total, trophectoderm and inner cell mass cell number, while vitrification lowered the inner cell mass proportion and increased apoptosis. Both exposures depolarised mitochondrial membrane potential and depleted glutathione; reactive oxygen species rose with an interaction, being highest in vitrified IVF embryos. IVF reduced live birth rate and litter size. In the adult offspring, high-resolution respirometry of isolated mitochondria from left ventricle revealed reduced oxidative phosphorylation capacity with an increased H_2_O_2_ production, altered OXPHOS subunit abundance and reduced complex I, III and IV activities.

**Conclusions:** IVF and vitrification impose distinct disturbance on preimplantation embryo redox states and bioenergetics, and this early disturbance is followed into adulthood with a reduced mitochondrial aerobic capacity and increased basal ROS production. These results have important implications for IVF practices and suggest that mitochondria may be permanently programmed by this procedure.

**Graphical Summary:** 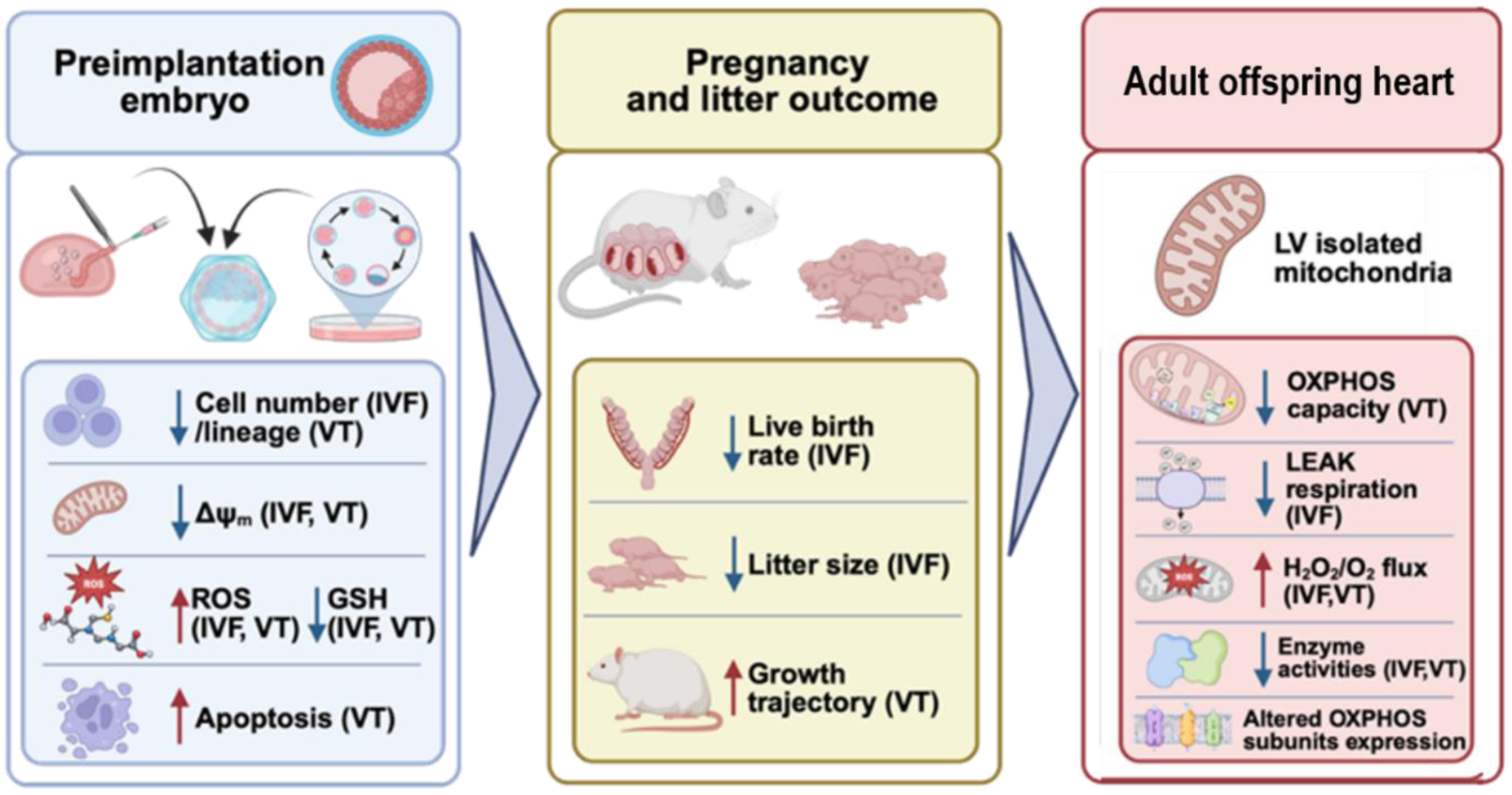

IVF and vitrification impose distinct and partly independent effects on the preimplantation embryo that persist into the adult offspring heart. At the blastocyst stage, IVF reduced cell number and vitrification altered lineage allocation, while both exposures lowered mitochondrial membrane potential (ΔΨm) and glutathione (GSH) and raised reactive oxygen species (ROS); vitrification additionally increased apoptosis. After embryo transfer, IVF reduced live birth rate and litter size, whereas vitrification altered postnatal growth trajectory. In adult offspring, ventricular mitochondria, vitrification reduced OXPHOS capacity and IVF reduced LEAK respiration, while both exposures increased H2O2/ O2 flux, reduced respiratory chain enzyme activities and altered OXPHOS subunit abundance.

## Introduction

In vitro fertilisation (IVF) has resulted in the birth of more than 10 million babies worldwide^1^, accounting for over 3% of overall births in the UK. Cryopreservation techniques such as vitrification are increasingly performed in clinics, which are now used in nearly half of UK treatment cycles^2^. These procedures coincide with a period of extensive embryonic epigenetic reprogramming^3^, and epidemiological studies have identified a broadly consistent cardiovascular phenotype in the resulting offspring. From fetal life onwards, IVF-conceived offspring show altered cardiac geometry and impaired myocardial function that may persist after birth into early childhood^4–6^, together with an increased incidence of congenital heart defects^7^. By school age this phenotype includes higher blood pressure, cardiac remodelling, and vascular and metabolic abnormalities^8–10^, and adolescence has been associated with premature vascular ageing and arterial hypertension^11^. These findings have also been supported by animal models, in which IVF-conceived offspring develop hypertension, endothelial dysfunction, altered left ventricular geometry and impaired glucose homeostasis, accompanied by epigenetic changes which regulate vascular and cardiac remodelling ^12–17^. This is particularly concerning because the periconceptional period is one of the earliest windows during which long-term physiological programming can occur^3,18^. However, evidence for these cardiac phenotypes that persist into human offspring adulthood remains limited and inconsistent ^19–22^, and almost all focuses on the cardiac structure and function rather than its bioenergetics, whether IVF and vitrification deposit a metabolic deficit on adult offspring heart has not been explored.

The heart is among the most oxidative tissues in the body, deriving approximately 90% of its ATP from mitochondrial oxidative phosphorylation (OXPHOS)^23^. Fetal heart relies on glycolysis and lactate oxidation as its energy source, but fatty acid oxidation (FAO) becomes dominant when the heart matures^24^. Mitochondria are also the principal source of reactive oxygen species (ROS), where a proportion of electrons slip from the respiratory chain at complex I and complex III and form superoxide^25^. Inefficient respiration and oxidant production are mechanistically coupled, and any developmental insult that compromises electron transfer would leave a heart in a less energetically capable and more redox imbalanced state. Adverse periconceptional conditions such as prenatal hypoxia and maternal undernutrition have been proved to programme persistent reductions in respiratory capacity, increased ROS production and altered respiratory complex activity in the adult offspring heart^26–30^. Critically, such defects can remain compensated at rest and emerge only under metabolic challenge^28^, and would therefore not to be discovered by structural and functional measures on which the current IVF literature has so far relied.

The preimplantation embryo is the most plausible point of origin for such a deficit. Its mitochondria are maternally inherited and morphologically immature, and they cannot replicate at early stages of development ^31^. IVF exposes the embryo to artificial media, altered substrate availability, repeated handling during precisely this period, and IVF-derived blastocysts show a Warburg-like shift towards glycolysis, elevated ROS and depleted glutathione^32^. Vitrification imposes a second and distinct insult through high cryoprotectant concentrations, osmotic shifts and ultra-rapid cooling and warming, which could cause mitochondrial DNA fragmentation and reduction in embryo respiration after warming ^33–35^. Whether this early redox lesion is transient or is propagated into the mitochondria of the adult heart remains unknown. Human studies cannot readily resolve this, because the effect of the procedure is confounded by the subfertility and advanced maternal age, and myocardial tissue is not available from otherwise healthy offspring. To our knowledge, only one animal study has examined cardiac mitochondria after assisted conception, reporting reduced myocardial mitochondrial abundance and lower OXPHOS complex I and IV protein in adolescent sheep after in vitro embryo culture^36^. Mitochondrial respiratory functions have never been measured in the heart of any IVF offspring, and the contribution of vitrification has never been separated from that of IVF, despite frozen embryo transfer now accounting for nearly half of UK cycles.

Here we used a mouse model to study the effects of IVF and vitrification on preimplantation embryo redox system and their long-term impact on adult offspring cardiac metabolism. We characterised the mitochondrial and redox phenotype of the blastocyst, then followed the IVF offspring to adulthood by assessing their cardiac structure and function via echocardiography and by measuring left ventricular mitochondrial respiration, ROS production, respiration chain complex abundance and activity. To our knowledge, this is the first direct measurement of cardiac mitochondrial function in the offspring of assisted conception of all studies.

## Materials and Methods

### Animals and ethical approval

All procedures were performed in accordance with the UK Home Office Animals (Scientific Procedures) Act 1986 and were approved by the local ethics committee at the University of Manchester (PD7C22AA9). CD-1 IGS mice (Charles River, Kent, UK) were maintained at the University of Manchester on a 07:00–19:00 light cycle with ad libitum access to food and water, and were acclimatised for at least one week before any procedure.

### Experimental design

Embryos and offspring were assigned to four groups defined by embryo origin and cryopreservation status: in vivo-fertilised blastocysts flushed from the uterus (FB), flushed blastocysts subjected to vitrification and warming (FB_VT), IVF-derived blastocysts (IVF), and IVF-derived blastocysts subjected to vitrification and warming (IVF_VT).

### Embryo collection, in vitro fertilisation and embryo culture

CD-1 females aged 4–6 weeks were superovulated with 5 IU pregnant mare’s serum gonadotrophin injection (PMSG; Intervet, UK) followed 48 h later by 5 IU human chorionic gonadotrophin injection (hCG; Chorulon, UK) and were immediately mated with stud males. Virginal plug was confirmed on the next morning (E0.5). For in vivo-fertilised blastocysts collection, female mice were sacrificed on E3.5 (90-96 h after hCG), blastocysts were flushed from uterus with M2 medium (MR-015-D, Merck) and transferred to KSOM medium (MR-106-D, Merck) under mineral oil (M5310, Merck) at 37 °C in 5% CO_2_ and 5% O_2_ atmosphere for culture or further experiments. For IVF, sperm from fertile CD-1 males was collected by puncturing the cauda epididymis and vas deferens with a needle in HTF (human tubal fluid medium; MR-070-D, Merck), and cumulus–oocyte complexes were retrieved from superovulated females 13–16 h after hCG. IVF was conducted in HTF at 37 °C for 4–6 h. Zygotes displaying two pronuclei and an extruded second polar body were cultured to the blastocyst stage in KSOM at 37 °C in an atmosphere containing 5% CO₂ and 5% O₂.

### Vitrification and warming

Blastocysts were cryopreserved with the Kitazato Cryotop open system (VT601 vitrification kit, VT602 warming kit and Cryotop devices; Kitazato, Japan) at room temperature in two steps: equilibration and vitrification. Briefly, up to six blastocysts were held in equilibration solution (ES) for 13 min. The vitrification step was completed within 90 s, during which the embryos were passed through two drops of vitrification solution (VS1 and VS2), loaded onto the tips of the Cryotops in minimal volume and plunged into liquid nitrogen. Devices were stored under liquid nitrogen for at least one week. Warming was initiated by transferring the Cryotop directly into thawing solution (TS) at 37 °C for 1 min, followed by diluent solution (DS) for 3 min and washing solutions WS1 and WS2 for 5 min and 1 min respectively. Warmed embryos were recovered for 4 h in KSOM; those failing to re-expand or showing fragmentation were classified as non-surviving and excluded.

### Embryo transfer

To induce pseudopregnancy, CD-1 females aged 8–10 weeks were mated with surgically vasectomised CD-1 males, and those with copulatory plugs the following morning were designated 0.5 days post-coitum (dpc). At 2.5 dpc, recipients were anaesthetised with isoflurane on a heated pad and the uterus was exteriorised through a flank incision. Under a stereomicroscope, the uterine wall was punctured and 14-15 blastocysts in M2 medium were expelled into single side of the uterine horn through a transfer pipette. After suturing, mice received buprenorphine (0.015 mg/kg, s.c.), were monitored until recovery from anaesthesia and were single-housed until parturition.

### Blastocyst immunostaining, lineage allocation and apoptosis

Blastocysts were cultured for a further 20 h to E4.5, washed three times in DPBS and fixed in 4% paraformaldehyde for 20 min at room temperature. All subsequent steps were performed in 1% w/v agarose-coated 96-well plates to prevent adhesion. Embryos were permeabilised in 0.55% v/v Triton X-100 in DPBS for 20 min and washed in DPBS-X (0.1% v/v Triton X-100 in DPBS). Apoptosis was detected before blocking using a one-step TUNEL assay (E-CK-A322, Stratech, UK): embryos were equilibrated for 15 min and incubated with terminal deoxynucleotidyl transferase and fluorescein-conjugated dUTP for 60 min at 37 °C, then washed three times in DPBS. Embryos were blocked in 10% v/v fetal bovine serum in DPBS-X for 1 h and incubated overnight at 4 °C with ready-to-use anti-CDX2 antibody (AM392-5M, Bio-Genex, USA), followed by goat anti-mouse IgG (H+L) Alexa Fluor 488 (1:200; A-11001, Invitrogen) for 1 h at room temperature. Nuclei were counterstained with Hoechst 33342 (250 µg/mL; B2261, Merck) for 15 min. Embryos were mounted in DPBS drops under mineral oil in glass-bottom dishes (P35G-1.5-14-C, MatTek, USA).

### Mitochondrial membrane potential, intracellular ROS and glutathione

All probes were applied to E3.5 blastocysts in KSOM for 30 min in the benchtop incubator. Mitochondrial membrane potential (ΔΨ_m_) was assessed live with the ratiometric probe JC-1 (10 µg/mL; T3168, Invitrogen), diluted fresh from a 10 mg/mL DMSO stock and centrifuged to remove undissolved dye, using 50 µM FCCP as a depolarised positive control; labelled blastocysts were washed in M2 and imaged immediately at 37 °C, with aggregate and monomer fluorescence reporting high and low ΔΨ_m_ respectively. Intracellular ROS and reduced glutathione were assessed with CellROX Green (5 µM; C10444, Invitrogen) and ThiolTracker Violet (20 µM; T10095, Invitrogen) in fixed blastocysts; blastocysts were washed in DPBS, fixed in 4% paraformaldehyde for 20 min and mounted in glass-bottom dishes under mineral oil, with CellROX-labelled embryos counterstained with Hoechst 33342 (250 µg/mL, 15 min) before mounting.

### Glucose uptake and ATP content

Both assays were performed on the individual blastocysts. Single E3.5 blastocysts were cultured in 2 µL KSOM microdrops under mineral oil for 24 h alongside cell-free control drops, after which the spent medium and the corresponding embryo were snap-frozen separately and stored at −80 °C for ATP measurement; dead or arrested embryos were excluded. Glucose was quantified by the NADPH-linked hexokinase/glucose-6-phosphate dehydrogenase fluorometric method ^37^, reading 1 µL of sample against 25 µL assay mix at 430/460 nm on a Synergy HTX reader (Agilent) before and 10 min after addition.

ATP was measured in the embryo lysate by luciferase bioluminescence (FLASC, Merck) . Both analytes were quantified against six-point standard curves (0–0.5 mM glucose; 0–10 nM ATP; curves with R² < 0.99 rejected) and expressed as pmol glucose per embryo per hour and fmol ATP per blastocyst.

### Offspring biometry and mitochondrial isolation

Litter size and sex ratio were recorded at birth, live birth rate was calculated as the number of pups born per embryo transferred (%), and body weight was recorded from 2 to 24 weeks of age.

Mitochondria were isolated by centrifugation in MSE buffer (5 mM MOPS, 2 mM Na-EDTA, 70 mM sucrose, 220 mM D-mannitol, pH 7.35 at 4 °C, with 0.1% m/v BSA added immediately before use). Following cervical dislocation the left ventricle was dissected, washed free of blood, minced, digested with 15.4 µM trypsin for 10 min and stopped with 10.6 µM trypsin inhibitor, then homogenised in a Potter–Elvehjem grinder (KIMBLE KONTES size 23, DWK) with a PTFE pestle at 135 rpm. The homogenate was centrifuged at 800 × *g*, and the filtered supernatant twice at 9000 × *g*; the final pellet was resuspended in 200 µL MSE and held on ice for immediate respirometry, with the remainder stored at −80 °C and immunoblotting aliquots supplemented with 1% v/v protease and phosphatase inhibitor cocktail. Mitochondrial protein was determined by Bradford assay (Quick Start, Bio-Rad) against a 0–5 mg/mL BSA curve at 595 nm, subtracting 1 mg/mL to account for BSA in MSE.

### High-resolution respirometry

Oxygen consumption and H_2_O_2_ production were measured simultaneously in an Oroboros Oxygraph-2k (Innsbruck, Austria). Measurements were in 2 mL chambers of MiR05 (0.5 mM EGTA, 3 mM MgCl_2_, 60 mM KMES, 20 mM taurine, 10 mM KH_2_PO_4_, 20 mM HEPES, 110 mM sucrose, pH 7.1 with 5 M KOH, 1 g/L BSA added after pH adjustment) at 37 °C. Horseradish peroxidase (1 U/mL), superoxide dismutase (5 U/mL) and Amplex Red (10 µM) were added before mitochondrial injection, and the H_2_O_2_ signal calibrated by two sequential 0.1 µM H₂O₂ additions.

For the carbohydrate SUIT protocol, malate (2 mM), pyruvate (5 mM) and glutamate (10 mM) were added to each chamber, followed by an isolated mitochondrial suspension (2.5 µL), to achieve the LEAK respiratory state through complex I in the absence of adenylates (LEAK_N,CI_). ADP (250 µM) was then added to activate OXPHOS with complex I substrates (OXPHOS_CI_), and respiration was left to occur until ADP was depleted to achieve the LEAK state in the presence of adenylates (LEAK_T,CI_). Succinate (10 mM) was then added (LEAK_T,CI+CII_), followed by a further ADP addition (250 µM) to measure both complex I and II substrates on OXPHOS (OXPHOS_CI+CII_). Mitochondria were then uncoupled through the addition of carbonyl cyanide-4-(trifluoromethoxy)phenylhydrazone (FCCP), in order to assess maximum electron transfer capacity (ET_CI+CII_); FCCP was titrated in 0.75 µM steps until no further increase in respiration rate. The complex I inhibitor rotenone (0.15 µM) was then added to assess ET_CII_. Next, antimycin A (12.5 µM), the complex III inhibitor, was added to block the electron transport pathway and measure residual nonmitochondrial O_2_ consumption (ROX). For complex IV activity determination, the electron donor N,N,N′,N′-tetramethyl-p-phenylenediamine (TMPD, 0.5 mM) was added, together with ascorbate (2 mM) to prevent TMPD auto-oxidation (ET_CIV_). Finally, the complex IV inhibitor sodium azide (50 mM) was added to measure background nonmitochondrial O_2_ consumption following TMPD addition. The respiratory control ratio (RCR) was calculated as OXPHOS_CI_/LEAK_T,CI_, and OXPHOS coupling efficiency as 1 − 1/RCR.

Fatty acid oxidation was assessed in a separate run. Diluted malate (0.08 mM) was added at a concentration sufficient to sustain NAD^+^ regeneration during β-oxidation without saturating the NADH-linked pathway, followed by an isolated mitochondrial suspension (3 µL) and palmitoyl-DL-carnitine (12.5 µM), to achieve the LEAK state with fatty acid substrate (LEAK_M+Pal_). ADP was then added to activate OXPHOS with fatty acid substrate (OXPHOS_M+Pal_), and palmitoyl-carnitine was titrated in 7.5 µM steps until additions failed to cause a subsequent increase in respiration rate. Values at each respiratory state were normalised to the protein concentration of the mitochondrial suspension, and H₂O₂ production was additionally expressed relative to O_2_ flux (H_2_O_2_/O_2_).

### Respiratory chain enzyme activities

Complex I–V activities were measured spectrophotometrically in isolated mitochondria on a Synergy HTX plate reader (Agilent), adapted from Spinazzi et al. and Brischigliaro et al. ^38,39^. V_max_ was derived in Gen5 software and activity expressed as µmol min^-1^ mg^-1^. Complex I: rotenone-sensitive (10 µM) NADH oxidation at 340 nm (ε = 6.2 mM^-1^ cm^-1^) in 50 mM potassium phosphate (KP) pH 7.5 with 3 mg/mL BSA, 300 µM KCN, 100 µM NADH, 80 µM DCPIP; initiated with 55.5 µM ubiquinone-1. Protein sample 80 µg/mL. Complex II: malonate-sensitive (10 mM) DCPIP reduction at 600 nm (ε = 19.1 mM^-1^ cm^-1^) in 25 mM KP pH 7.5 with 1 mg/mL BSA, 20 mM succinate, 300 µM KCN, 80 µM DCPIP; initiated with 46.3 µM decylubiquinone. Protein sample 48 µg/mL. Complex III: antimycin A-sensitive (18.5 µM) cytochrome c reduction at 550 nm (ε = 18.5 mM^-1^ cm^-1^) in 25 mM KP pH 7.5 with 0.025% Tween-20, 100 µM EDTA, 500 µM KCN, 75 µM cytochrome c; initiated with 92.6 µM decylubiquinol. Sample 2 µg/mL. Complex IV: KCN-sensitive (300 µM) oxidation of reduced cytochrome c at 550 nm (ε = 18.5 mM^-1^ cm^-1^) in 50 mM KP pH 7.0; initiated with 0.37 mM reduced cytochrome c. Sample 4.8 µg/mL. Complex V (assayed as ATP hydrolysis): oligomycin-sensitive (2 µg/mL) NADH oxidation coupled through pyruvate kinase and lactate dehydrogenase at 340 nm (ε = 6.2 mM^-1^ cm^-1^) in 25 mM KP pH 7.2 with 5 mM MgCl_2_, 100 mM KCl, 2.5 mg/mL BSA, 2 mM phosphoenolpyruvate, 0.2 mM NADH, 3 U/mL pyruvate kinase, 3 U/mL lactate dehydrogenase; initiated with 4.63 mM ATP. Sample 68 µg/mL.

### Immunoblotting of OXPHOS subunits

Isolated mitochondria (5 µg protein) in NuPAGE sample buffer with reducing agent were denatured at 50 °C for 10 min, proteins were separated on 4–12% NuPAGE Bis-Tris midi gels in MES buffer (60 V for 10 min, then 200 V for 40 min, on ice) and transferred to 0.45 µm nitrocellulose (Trans-Blot Turbo, Bio-Rad; 7 min). Membranes were stained with Revert 700 Total Protein Stain (LI-COR) for lane normalisation, blocked in 5% w/v milk in TBS-T for 1 h, and incubated overnight at 4 °C with Total OXPHOS Rodent WB Antibody Cocktail (1:1000; ab110413, Abcam) detecting NDUFB8 (complex I), SDHB (complex II), UQCRC2 (complex III), MTCO1 (complex IV) and ATP5A (complex V), followed by IRDye 800CW donkey anti-mouse IgG (1:20,000; LI-COR) for 1 h at room temperature. Membranes were imaged on a LI-COR Odyssey CLx, and band intensities were quantified in Image Studio and normalised to total protein within each lane and to a common internal control sample across membranes.

### Confocal imaging and image analysis

Images were acquired on a Leica SP8x confocal microscope (1 Airy unit pinhole, 400 Hz scan speed, 512 × 512 pixels, sequential channel acquisition; LAS-X software). Lineage allocation was counted in Imaris 10 (Oxford Instruments, UK) by manual annotation of CDX2-positive (trophectoderm) and Hoechst-positive/CDX2-negative (inner cell mass) nuclei across reconstructed z-stacks. Remaining analyses used ImageJ on summed z-stacks with background subtraction: TUNEL signal was thresholded and expressed as a fraction of total Hoechst-stained nuclear area; JC-1 as the aggregate:monomer ratio normalised to the FCCP control of the same replicate; and CellROX and ThiolTracker as total fluorescence per embryo area, with ThiolTracker additionally normalised to the FB group.

### Statistical analysis

Analyses were performed in GraphPad Prism v11.0.0, SPSS Statistics (IBM) and RStudio (Posit Software). Normality was assessed by Shapiro–Wilk test and quantile–quantile plots, and homogeneity of variance by Levene’s test. Data were analysed with embryo type (FB vs IVF) and cryopreservation (fresh vs vitrified–warmed) as fixed factors: datasets meeting parametric assumptions by two-way ANOVA, and those with heterogeneous variance by generalised least squares (GLS) models. TUNEL-positive area fraction, which contained excess zeros, was analysed by generalised linear model with a Tweedie distribution and log link. Echocardiographic functional parameters were analysed by ANCOVA (or GLS ANCOVA where appropriate) with heart rate as a covariate and are reported as estimated marginal means evaluated at the mean heart rate. Šidák-adjusted post hoc comparisons were applied where an interaction was significant. Significance was set at P < 0.05, denoted *P < 0.05, **P < 0.01, ***P < 0.001, ****P < 0.0001.

## Results

### IVF reduces blastocyst cell number, whereas vitrification alters lineage allocation and increases apoptosis

To determine whether IVF and vitrification affected blastocyst quality, embryos from all four groups were cultured to E4.5 and stained for total nuclei (Hoechst), the trophectoderm marker CDX2, and apoptotic nuclei (TUNEL) (Figure 1A).

**Fig 1.**
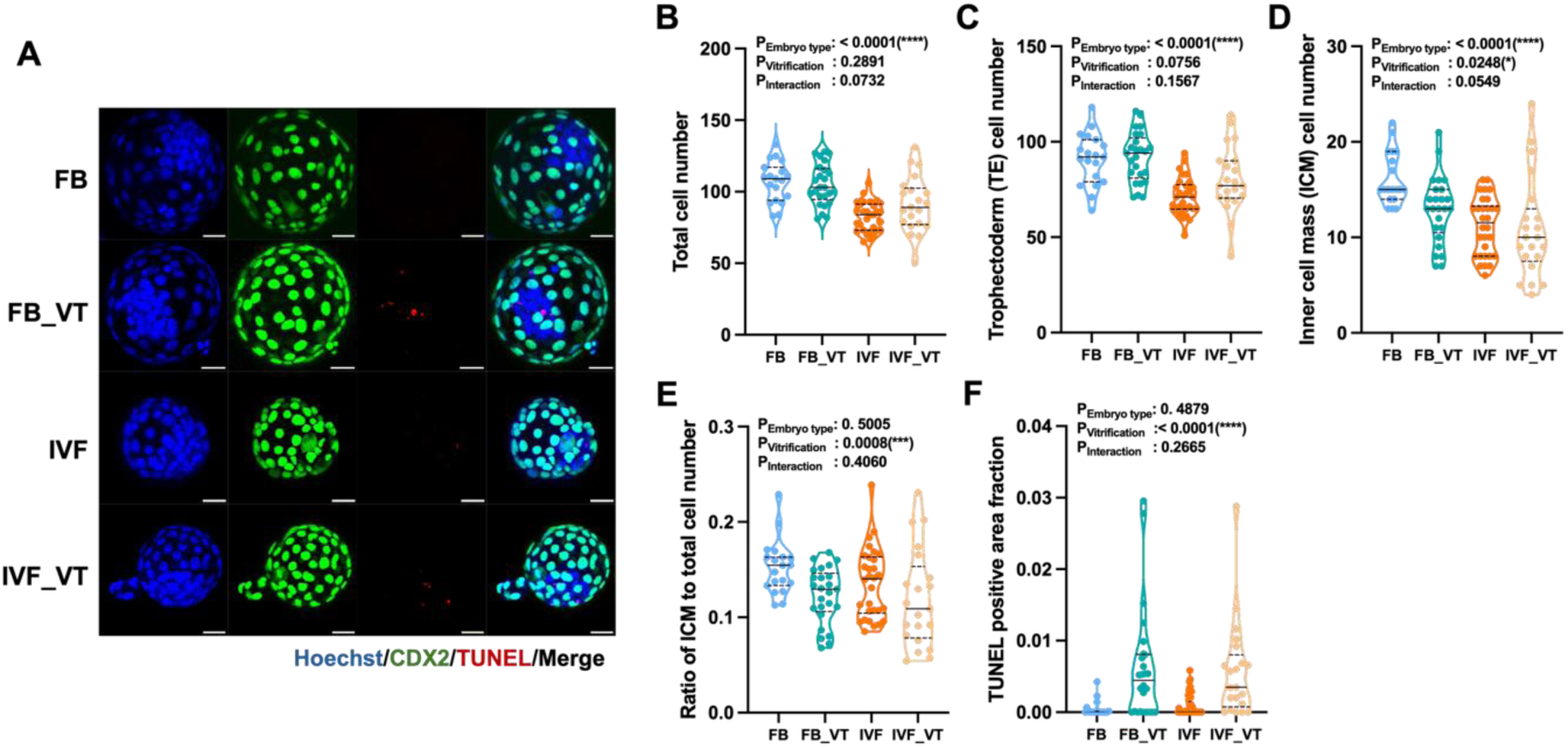
IVF reduces blastocyst cell number, whereas vitrification alters lineage allocation and increases apoptosis. **(A)** Representative immunofluorescence images of E4.5 blastocysts from each group stained for total nuclei (Hoechst, blue), the trophectoderm marker CDX2 (green) and apoptotic nuclei (TUNEL, red), with merged images. Scale bar: 25 µm. **(B)** Total cell number. **(C)** Trophectoderm (TE) cell number. **(D)** Inner cell mass (ICM) cell number, calculated by subtracting TE cell number from total cell number. **(E)** Ratio of ICM to total cell number. **(F)** TUNEL-positive area expressed as a fraction of total Hoechst-stained nuclear area. Data are violin plots showing individual embryos, with solid lines indicating the median and dashed lines the quartiles. N = 19 (FB), 25 (FB_VT), 31 (IVF), 21 (IVF_VT). **B–D** were analysed by two-way ANOVA; **E** by a generalised least squares (GLS) model owing to heteroscedasticity; and **F** by a generalised linear model with a Tweedie distribution and log link owing to excess zeros. Main effect and interaction *P* values are shown within each panel. No interaction reached significance, so no post hoc comparisons were performed.

IVF reduced total cell number (*P* < 0.0001), trophectoderm (TE) cell number (*P* < 0.0001) and inner cell mass (ICM) cell number (*P* < 0.0001), with no significant interaction in any measure (Figure 1B–D). Vitrification additionally reduced ICM cell number (*P* = 0.0248) but did not affect total or TE cell number. Consequently, the ICM:total cell ratio was reduced by vitrification (*P* = 0.0008) with no effect of embryo type (*P* = 0.5005), indicating a shift in lineage allocation in favour of the trophectoderm that was independent of how the embryo was fertilised (Figure 1E). Vitrification also increased the TUNEL-positive area fraction (*P* < 0.0001), again independently of embryo type (*P* = 0.4879) (Figure 1F).

IVF therefore constrained overall proliferative expansion, whereas vitrification acted selectively on the ICM and on apoptosis. Notably, IVF did not increase apoptosis, indicating that its effect on cell number was not attributable to cell death at the time of assessment.

### IVF and vitrification disturb blastocyst mitochondrial redox homeostasis and metabolism

Mitochondrial membrane potential (ΔΨ_m_), intracellular ROS, glutathione (GSH), glucose uptake and ATP content were measured in E3.5 blastocysts (Figure 2). JC-1 imaging showed a strong aggregate signal in FB blastocysts and progressively weaker aggregate relative to monomer fluorescence in FB_VT, IVF and IVF_VT embryos, with FCCP-treated embryos confirming probe responsiveness (Figure 2A). Both IVF (*P* < 0.0001) and vitrification (*P* < 0.0001) reduced the normalised aggregate:monomer ratio, with no interaction (*P* = 0.3179), indicating that the two exposures depolarised mitochondria independently and additively (Figure 2B).

**Fig 2.**
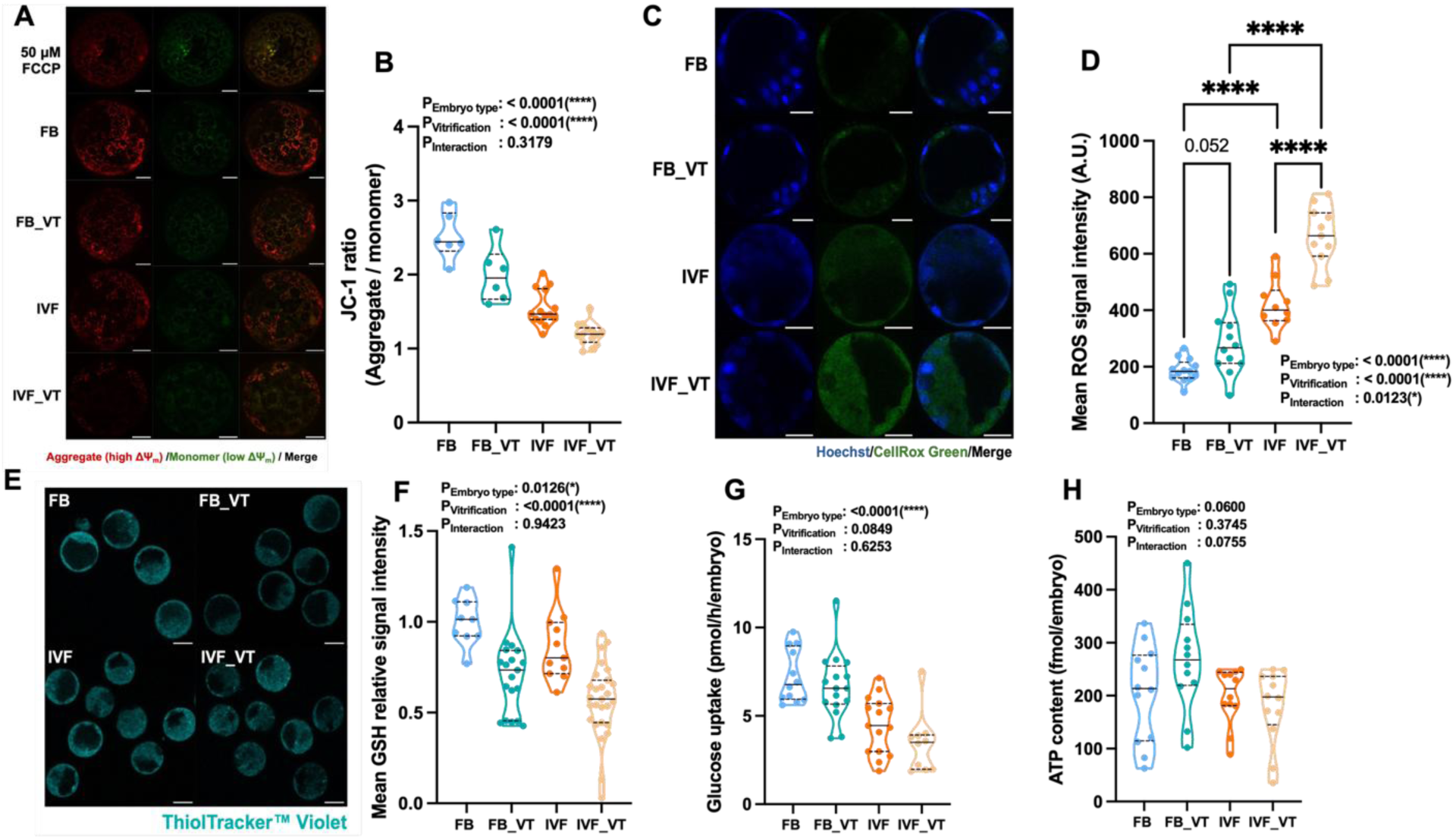
IVF and vitrification independently reduce blastocyst mitochondrial membrane potential and glutathione and increase ROS, with IVF-derived embryos more susceptible to vitrification-induced oxidative stress. **(A)** Representative JC-1 images of live E3.5 blastocysts showing aggregate (red, high ΔΨm), monomer (green, low ΔΨm) and merged fluorescence. Embryos treated with 50 µM FCCP are shown as a depolarised control. Scale bar: 25 µm **(B)** JC-1 aggregate:monomer ratio, normalised to the FCCP control of the same replicate. N = 6 (FB), 6 (FB_VT), 14 (IVF), 14 (IVF_VT). **(C)** Representative CellROX Green images with Hoechst counterstain (blue) and merged fluorescence. Scale bar: 25 µm **(D)** Mean intracellular ROS signal intensity. N = 13 (FB), 12 (FB_VT), 10 (IVF), 11 (IVF_VT). **(E)** Representative ThiolTracker Violet images of the four groups. Scale bar: 25 µm **(F)** Mean glutathione signal intensity, normalised to the FB group. N = 10 (FB), 19 (FB_VT), 11 (IVF), 27 (IVF_VT). **(G)** Glucose uptake per blastocyst per hour. N = 12 (FB), 16 (FB_VT), 15 (IVF), 10 (IVF_VT). **(H)** ATP content per blastocyst. N = 12 (FB), 12 (FB_VT), 11 (IVF),12 (IVF_VT). **G** and **H** were analysed by two-way ANOVA; **B, D** and **F** by GLS models owing to heteroscedasticity. Where the interaction was significant, Šidák-adjusted post hoc comparisons were applied. Main effect and interaction *P* values are shown within each panel. \**P* < 0.05, \*\**P* < 0.01, \*\*\**P* < 0.001, \*\*\*\**P* < 0.0001.

Intracellular ROS were increased by both IVF (*P* < 0.0001) and vitrification (*P* < 0.0001), and here the two factors interacted (*P* = 0.0123) (Figure 2C, D). Post hoc comparison showed that vitrification raised ROS markedly in IVF-derived blastocysts (IVF vs IVF_VT, *P* < 0.0001) while the same comparison in flushed blastocysts did not reach significance (FB vs FB_VT, *P* = 0.052). IVF-derived embryos were therefore less able to tolerate the additional oxidative burden of cryopreservation. Reduced glutathione was lowered by both IVF (*P* = 0.0126) and vitrification (*P* < 0.0001) without interaction (*P* = 0.9423) (Figure 2E, F), so the rise in ROS occurred alongside a depleted thiol buffer rather than being offset by it.

Glucose uptake was reduced by IVF (*P* < 0.0001) with no effect of vitrification (*P* = 0.0849) (Figure 2G). ATP content did not differ significantly between groups (Figure 2H). Blastocyst ATP content was maintained despite mitochondrial and redox disturbance.

### IVF reduces litter size and live birth rate, whereas vitrification increases adult body weight

Live birth rate (*P* < 0.0001) and litter size (*P* < 0.0001) were both reduced by IVF, with no effect of vitrification and no interaction (Figure 3A, B). Offspring sex ratio was unaffected (Figure 3C). Because the FB control group also underwent blastocyst recovery, handling and embryo transfer, these reductions are attributable to the IVF procedure rather than to superovulation, transfer or the recipient uterine environment.

**Fig 3.**
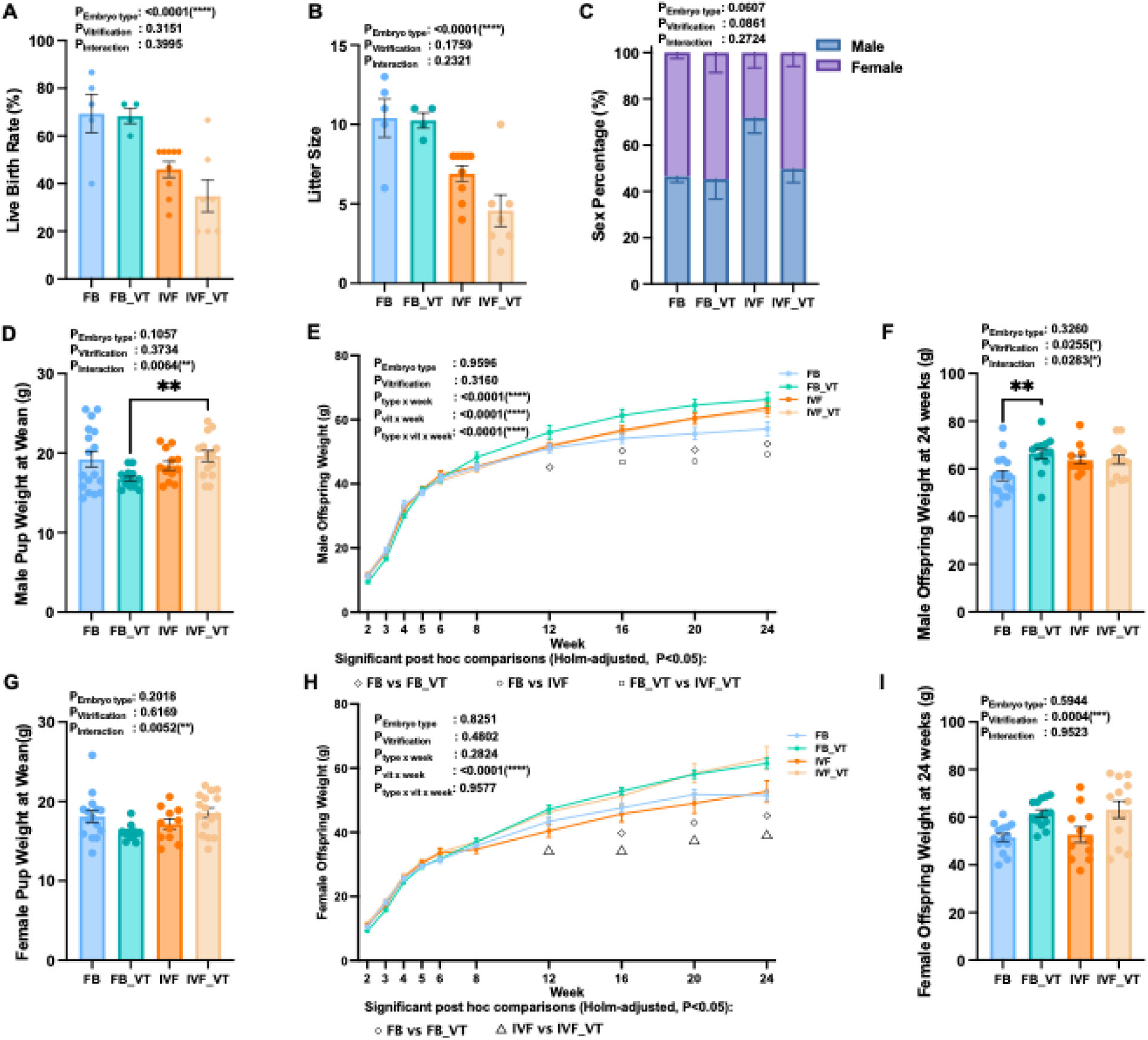
IVF reduces litter size and live birth rate, whereas vitrification increases adult body weight in both sexes. **(A)** Live birth rate, calculated as the number of pups born per embryo transferred. **(B)** Litter size. **(C)** Offspring sex distribution. **(D)** Male body weight at weaning. **(E)** Male body weight from 2 to 24 weeks of age. **(F)** Male body weight at 24 weeks. **(G)** Female body weight at weaning. **(H)** Female body weight from 2 to 24 weeks of age. **(I)** Female body weight at 24 weeks. Data are mean ± SEM with individual values shown; C is presented as stacked proportions. N = 4-9 litters per group for A–C and 11-17 offspring per group for D–I. **A, B, D, F, G** and **I** were analysed by two-way ANOVA with embryo type and vitrification as fixed factors. Growth trajectories **E, H** were analysed by three-way repeated-measures ANOVA with embryo type, vitrification and week as factors, followed by Holm-adjusted post hoc comparisons at each timepoint; significant pairwise comparisons (*P* < 0.05) are denoted ◇ (FB vs FB_VT), □ (FB vs IVF), ○ (FB_VT vs IVF_VT) and △ (IVF vs IVF_VT). Main effect and interaction *P* values are shown within each panel. \**P* < 0.05, \*\**P* < 0.01, \*\*\**P* < 0.001, \*\*\*\**P* < 0.0001.

Weaning weight showed no main effects in either sex but significant interactions in males (*P* = 0.0064) and females (*P* = 0.0052) (Figure 3D, G); in males, vitrification lowered weaning weight in the FB lineage but raised it in the IVF lineage. Growth trajectories diverged with age in both sexes (vitrification × week, *P* < 0.0001 for both; Figure 3E, H), with an additional three-way interaction in males (*P* < 0.0001), indicating that the effect of vitrification depended on both embryo type and age: curves were superimposable to week 8, after which vitrification progressively raised body weight in the FB lineage (FB vs FB_VT significant at weeks 12–24) while producing no clear separation within the IVF lineage, and IVF alone raised weight relative to FB only at weeks 20 and 24. By 24 weeks, vitrification had increased body weight in males (*P* = 0.0255, interaction *P* = 0.0283, confined within the FB lineage) and in females (*P* = 0.0004) (Figure 3F, I). The effect of vitrification on adult body weight was therefore consistent in females and lineage-dependent in males.

### Vitrification suppresses coupled and maximal respiration, whereas IVF lowers LEAK and fatty acid-supported respiration

Mitochondrial respiration was measured in isolated left ventricular mitochondria by high-resolution respirometry using a carbohydrate SUIT protocol (Figure 5A).

**Fig 5.**
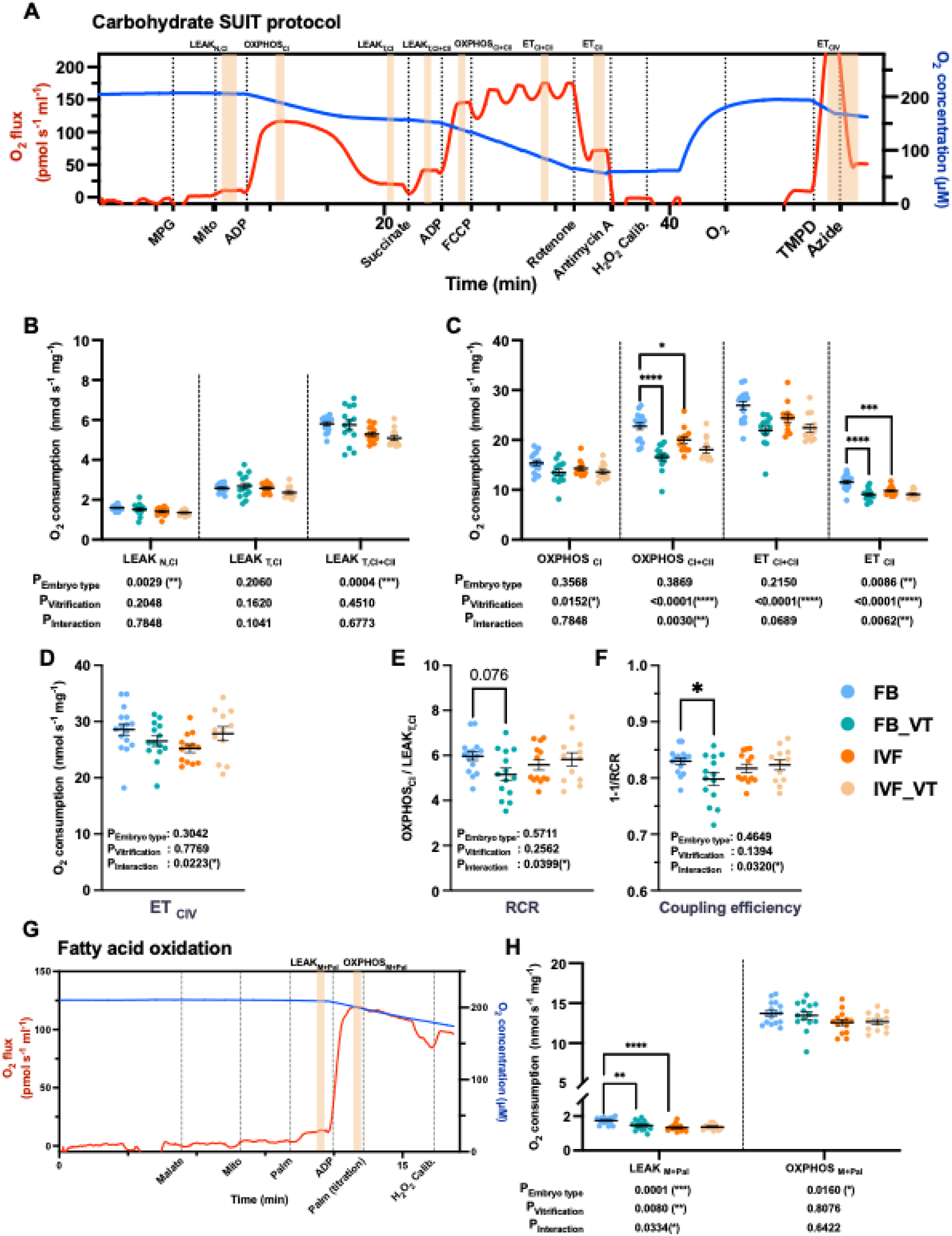
Vitrification suppresses coupled and maximal respiration, whereas IVF lowers LEAK and fatty acid-supported respiration in offspring left ventricular mitochondria. **(A)** Representative high-resolution respirometry trace for the carbohydrate SUIT protocol, showing oxygen flux (red) and oxygen concentration (blue) with sequential substrate and inhibitor additions indicated. Shaded bands denote the respiratory states quantified in **B–D**. **(B)** LEAK respiration with NADH-linked substrates in the absence of adenylates (LEAKN,CI), in the presence of adenylates (LEAKT,CI), and after addition of succinate (LEAKT,CI+CII). **(C)** Coupled respiration supported by complex I (OXPHOSCI) and by convergent complex I and II input (OXPHOSCI+CII), and maximal noncoupled electron transfer capacity with both substrate combinations (ETCI+CII) and with complex II substrates alone after complex I inhibition (ETCII). **(D)** Maximal complex IV capacity assessed with TMPD as artificial electron donor (ETCIV). **(E)** Respiratory control ratio (RCR), calculated as OXPHOSCI / LEAKT,CI. **(F)** OXPHOS coupling efficiency, calculated as 1 − 1/RCR. **(G)** Representative trace for the fatty acid oxidation protocol. **(H)** LEAK and coupled respiration supported by palmitoyl-carnitine (LEAKM+Pal, OXPHOSM+Pal). Oxygen flux is normalised to mitochondrial protein. Data are individual biological replicates with mean ± SEM; n = 12–16 per group. Analysis was by two-way ANOVA followed by Šidák-adjusted post hoc comparisons, except LEAKT,CI which was analysed by a GLS model owing to heteroscedasticity. Main effect and interaction *P* values are shown beneath each panel. \**P* < 0.05, \*\**P* < 0.01, \*\*\**P* < 0.001, \*\*\*\**P* < 0.0001.

IVF reduced LEAK respiration in the absence of adenylates (LEAK_N,CI_, *P* = 0.0029) and with convergent complex I and II input (LEAK_T,CI+CII_, *P* = 0.0004), with no effect of vitrification on either (Figure 5B). LEAK_T,CI_ was unchanged.

The coupled and maximal states showed the opposite pattern. Vitrification reduced OXPHOS_CI_ (*P* = 0.0152), OXPHOS_CI+CII_ (*P* < 0.0001), maximal noncoupled electron transfer capacity ET_CI+CII_ (*P* < 0.0001) and complex II-supported ET_CII_ (*P* < 0.0001) (Figure 5C). IVF additionally reduced ET_CII_ (*P* = 0.0086). Significant interactions for OXPHOS_CI+CII_ (*P* = 0.0030) and ET_CII_ (*P* = 0.0062) indicated that each procedure reduced these states only in the absence of the other: IVF was effective in fresh offspring and vitrification in the FB lineage, with no further reduction when both were applied. ET_CIV_ showed no main effects but a significant interaction (*P* = 0.0223) with no significant post hoc difference (Figure 5D).

Neither the respiratory control ratio nor OXPHOS coupling efficiency showed a main effect, but both showed significant interactions (*P* = 0.0399 and *P* = 0.0320), with vitrification reducing coupling efficiency in FB offspring only (Figure 5E, F).

Under fatty acid-supported conditions (Figure 5G), LEAK_M+Pal_ was reduced by both IVF (*P* = 0.0001) and vitrification (*P* = 0.0080) with a significant interaction (*P* = 0.0334) (Figure 5H): FB offspring had the highest fatty acid-supported LEAK respiration, which was lowered both by vitrification within the FB lineage (FB vs FB_VT, *P* < 0.01) and by IVF among fresh offspring (FB vs IVF, *P* < 0.0001), whereas IVF and IVF_VT offspring were indistinguishable. OXPHOS_M+Pal_ was reduced by IVF alone (*P* = 0.0160). The reduction of OXPHOS_M+Pal_ in IVF offspring reflects a lower intrinsic capacity to oxidise fatty acid-derived substrate.

### Both IVF and vitrification increase mitochondrial H₂O₂ production, with the increase persisting after complex I inhibition

H_2_O_2_ production was recorded simultaneously with oxygen flux and normalised to oxygen consumption for each respiratory state (Figure 6A).

**Fig 6.**
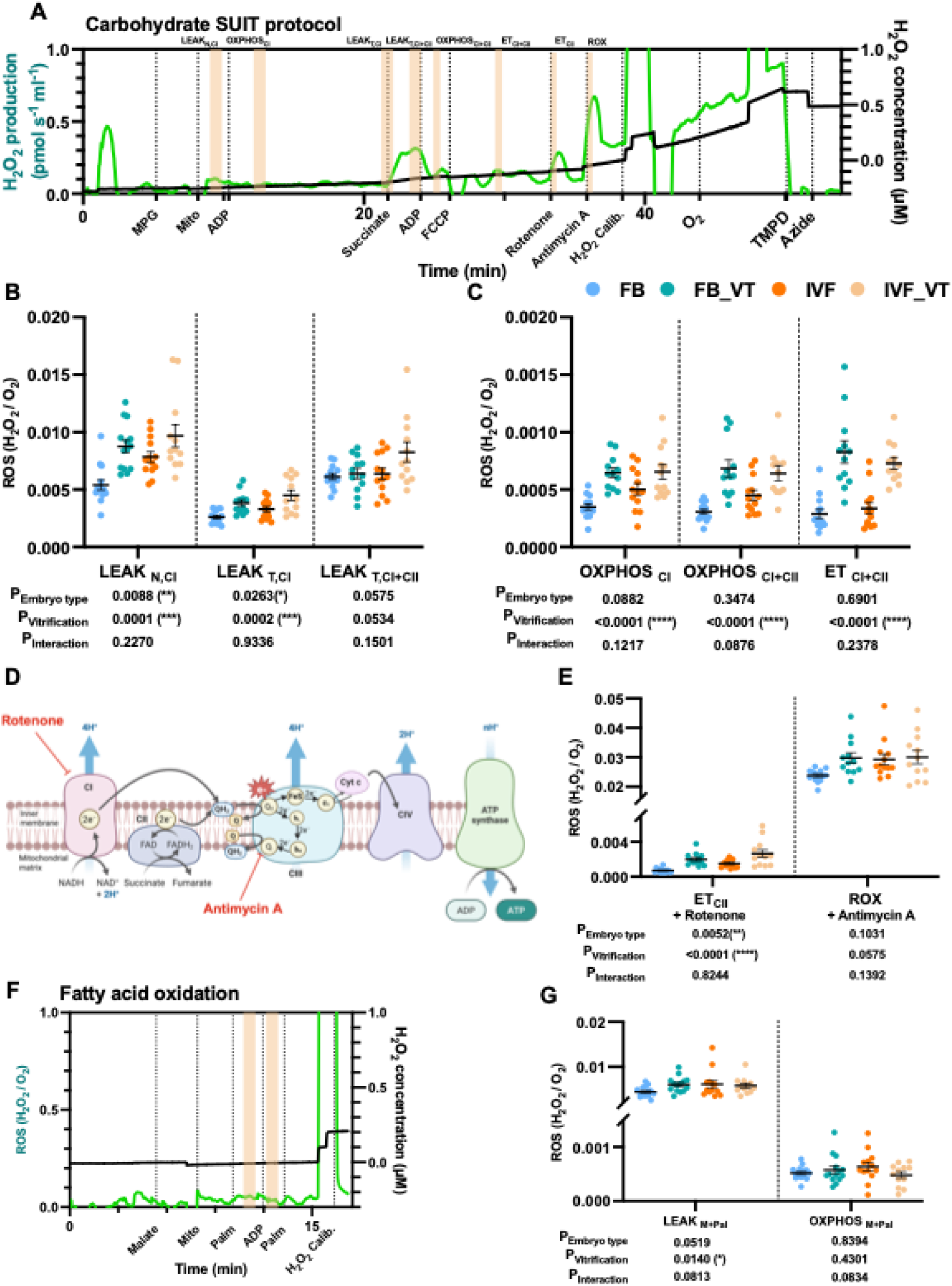
IVF and vitrification increase mitochondrial H_2_O_2_ production across multiple respiratory states, with the increase persisting after complex I inhibition and lost after complex III inhibition. **(A)** Representative trace of H2O2 production (green) and H2O2 concentration (black) recorded simultaneously with oxygen flux during the carbohydrate SUIT protocol. **(B)** H₂O₂ production normalised to oxygen flux (H2O2/O2) in the LEAK states (LEAKN,CI, LEAKT,CI, LEAKT,CI+CII). **(C)** H2O2/O2 in the coupled and noncoupled states (OXPHOSCI, OXPHOSCI+CII, ETCI+CII). **(D)** Schematic of the electron transport chain indicating inhibition of complex I by rotenone and of the complex III Qi site by antimycin A (Created with BioRender.com). **(E)** H₂O₂/O₂ after complex I inhibition with rotenone (ETCII) and after complex III inhibition with antimycin A (ROX). **(F)** Representative H2O2 trace for the fatty acid oxidation protocol. **(G)** H2O2/O2 during palmitoyl-carnitine-supported LEAK and coupled respiration (LEAKM+Pal, OXPHOSM+Pal). Data are individual biological replicates with mean ± SEM; n = 12-16 per group. Analysis was by two-way ANOVA. Main effect and interaction *P* values are shown beneath each panel. No interaction reached significance in any state.

Both IVF and vitrification increased H_2_O_2_/O_2_ flux in the complex I-supported LEAK states (LEAK_N,CI_: *P* = 0.0088 and *P* = 0.0001; LEAK_T,CI_: *P* = 0.0263 and *P* = 0.0002) (Figure 6B). Vitrification independently increased H_2_O_2_/O_2_ during OXPHOS_CI_ (*P* < 0.0001), OXPHOS_CI+CII_ (*P* < 0.0001) and ET_CI+CII_ (*P* < 0.0001) (Figure 6C). No interaction was significant in any state, indicating that the two exposures acted independently.

Two pharmacological observations could localise the source of the excess H_2_O_2_ (Figure 6D). The between-group differences persisted after complex I inhibition with rotenone, where both IVF (P = 0.0052) and vitrification (P < 0.0001) increased H_2_O_2_/O_2_ in the ET_CII_ state (Figure 6E), indicating that the phenotype does not depend predominantly on complex I, including reverse electron transport through its Q-binding site. The differences were then abolished after complex III inhibition with antimycin A (ROX; embryo type P = 0.1031, vitrification P = 0.0575) (Figure 6E). Antimycin A blocks the Q_i_ site and, by preventing normal Q-cycle turnover, this imposes a maximal Q_o_-site ROS-producing state; convergence under this condition therefore indicates that the phenotype depends on electron transfer through complex III. This indicates that the increased H_2_O_2_ most plausibly originates at the Q_o_ site of complex III, where ubiquinol oxidation generates a ubisemiquinone intermediate able to reduce O_2_ directly to superoxide. Complex II can generate ROS at its flavin site **under particular conditions**, but is a more conditional source and is not specifically implicated by this pharmacological profile.

Under fatty acid-supported conditions, vitrification increased H_2_O_2_/O_2_ during LEAK_M+Pal_ (P = 0.0140) but not during OXPHOS_M+Pal_ (Figure 6F, G).

### OXPHOS subunit abundance and catalytic activity dissociate, converging on complex III

To determine whether the respiratory phenotype reflected loss of respiratory chain protein, abundance of a representative subunit of each complex was measured by immunoblotting against total protein (Figure 7A, B).

**Fig 7.**
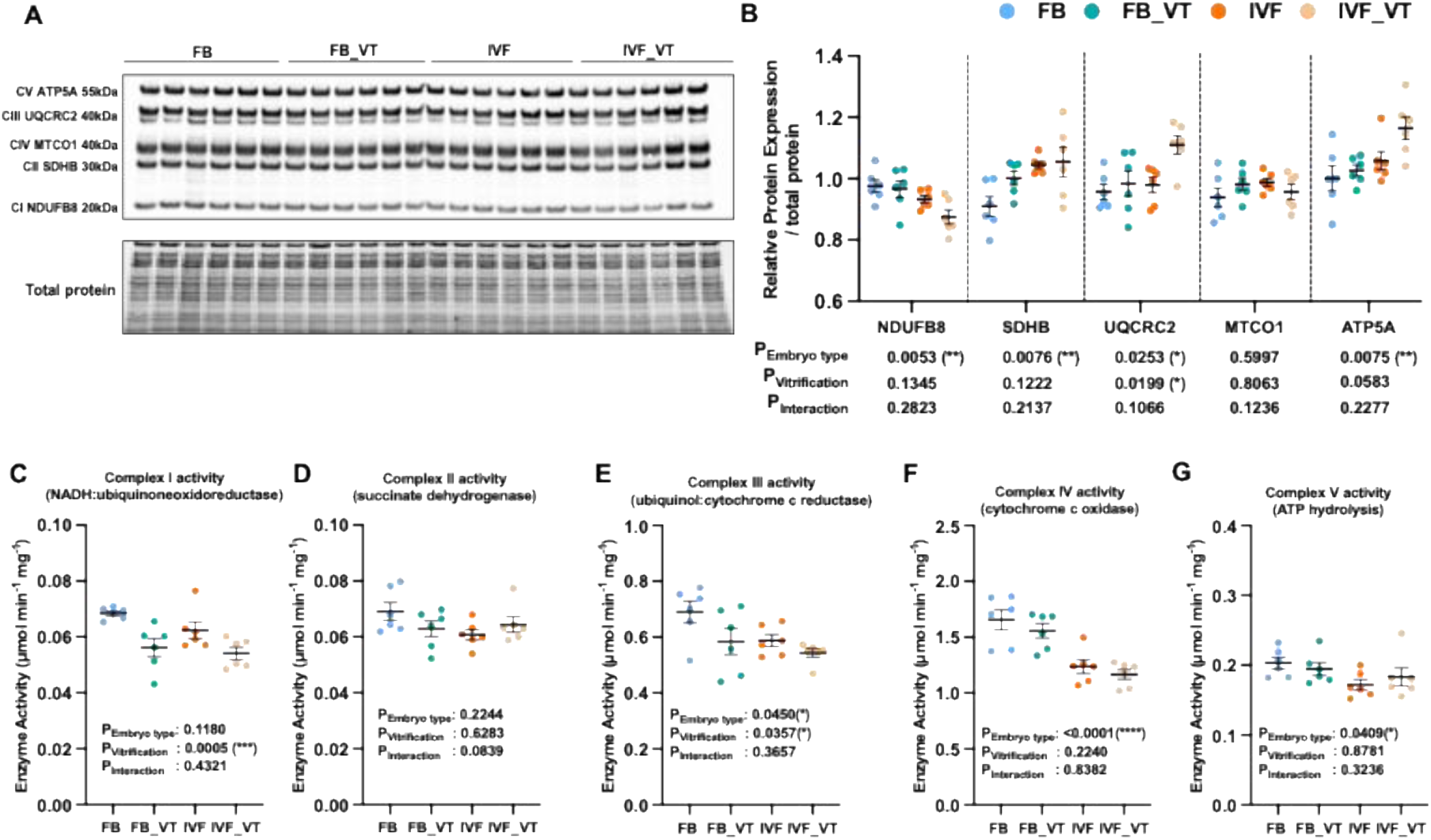
OXPHOS subunit abundance and respiratory chain enzyme activity dissociate in offspring left ventricular mitochondria, converging on complex III. **(A)** Representative immunoblot of five OXPHOS subunits detected with the Total OXPHOS Rodent antibody cocktail: NDUFB8 (complex I, 20 kDa), SDHB (complex II, 30 kDa), UQCRC2 (complex III, 48 kDa), MTCO1 (complex IV, 40 kDa) and ATP5A (complex V, 55 kDa). Total protein stain is shown below as the lane-normalisation control. **(B)** Relative abundance of each subunit normalised to total protein within each lane and to a common internal control across membranes. **(C–G)** Catalytic activity of **(C)** complex I (NADH:ubiquinone oxidoreductase), **(D)** complex II (succinate dehydrogenase), **(E)** complex III (ubiquinol:cytochrome c reductase), **(F)** complex IV (cytochrome c oxidase) and **(G)** complex V (assayed as ATP hydrolysis), each measured spectrophotometrically as the inhibitor-sensitive rate and expressed per mg mitochondrial protein. Data are individual biological replicates with mean ± SEM; n = 6 per group. Analysis was by two-way ANOVA. Main effect and interaction P values are shown within each panel. No interaction reached significance.

Changes were selective and bidirectional rather than uniform. IVF reduced NDUFB8 (complex I, *P* = 0.0053) but increased SDHB (complex II, *P* = 0.0076), UQCRC2 (complex III, *P* = 0.0253) and ATP5A (complex V, *P* = 0.0075). Vitrification independently increased UQCRC2 (*P* = 0.0199). MTCO1 (complex IV) was unchanged. Because all measurements were normalised to mitochondrial protein, these differences are unlikely to reflect a change in mitochondrial content.

Catalytic activities did not follow subunit abundance (Figure 7C–G). Complex I activity was reduced by vitrification alone (*P* = 0.0005). Complex II activity was unchanged. Complex III activity was reduced by both IVF (*P* = 0.0450) and vitrification (*P* = 0.0357). Complex IV (*P* < 0.0001) and complex V (*P* = 0.0409) activities were reduced by IVF alone.

Complex III was therefore the only complex whose activity was reduced by both procedures, and it was reduced despite an increase in UQCRC2 abundance. The same dissociation applied to complex V, where increased ATP5A abundance accompanied reduced activity. Together these data indicate selective remodelling of the respiratory chain with incomplete compensation, rather than a simple loss of OXPHOS protein.

## Discussion

Our study shows, for the first time, that cardiac mitochondrial respiratory function and ROS production are altered in the adult offspring of assisted conception. Specifically, we found that IVF and vitrification each disturbed blastocyst mitochondrial membrane potential, intracellular ROS and glutathione in embryos that had reached the blastocyst stage and were competent to establish pregnancy, and that mitochondria from the adult left ventricle had reduced respiratory capacity and elevated ROS production. The two procedures acted on largely separate parts of the electron transport system but converged functionally on complex III. Taken together, these results indicate that IVF and cryopreservation alter mitochondrial metabolism through largely independent routes, and these problems extend into adulthood. These findings have implications for embryo selection, and for the growing proportion of ART-conceived children exposed to both procedures.

### IVF and vitrification disturb blastocyst mitochondrial and redox homeostasis through distinct routes

The mitochondrial and redox disturbance we observed was present in blastocysts that went on to be transferred and establish pregnancies. Because blastocyst formation remains a principal criterion for clinical embryo selection^40–42^, this dissociation matters: reaching the blastocyst stage did not indicate that mitochondrial and redox homeostasis were intact.

IVF constrained proliferative expansion and reduced total and trophectoderm cell number without increasing apoptosis, whereas vitrification acted selectively on the inner cell mass, shifted lineage allocation towards the trophectoderm and increased TUNEL-positive area. Because apoptosis was not increased by IVF, the reduction in cell number more likely reflects fewer completed cell divisions than the loss of blastomeres already formed.

The significant interaction for intracellular ROS indicates that IVF leaves the embryo less able to tolerate a subsequent oxidative insult. This aligns with the finding that in vivo-derived blastocysts showed lower apoptosis and greater cryotolerance than in vitro-produced ones in bovine blastocysts^43^. Because glutathione was simultaneously depleted in our model, the rise in ROS is more likely to reflect both increased generation and diminished thiol buffering than either alone.

ATP content was preserved despite the fall in membrane potential. This is not the expected result: the proton-motive force reported by JC-1 is what drives ATP synthase, so a depolarised mitochondrial population would be expected to produce less ATP. Three explanations are possible. The first is compensatory glycolysis, which yields ATP independently of membrane potential and has been reported in IVF-generated murine blastocysts together with reduced mitochondrial respiration and depleted glutathione^32^. Our data do not support it, since glucose uptake was reduced rather than increased by IVF. The second is that energetic demand fell in parallel with supply, leaving the steady-state pool unchanged; lower glucose uptake alongside maintained ATP is consistent with this, and with the quiet embryo hypothesis, in which viable embryos operate at lower rather than higher metabolic turnover ^44,45^. The third is methodological: ATP content is a static pool measurement, and a pool can be held constant even when turnover falls, so preserved content does not establish preserved ATP production. Direct flux measurement would be required to distinguish these.

### Vitrification, not IVF, drives offspring growth, mirroring the human frozen embryo transfer phenotype

The two procedures also separated at the level of the whole animal. IVF reduced litter size and live birth rate without affecting adult body weight, whereas vitrification left litter outcomes intact but was the dominant influence on adult weight, increasing it uniformly in females and in an embryo origin-dependent manner in males. Because the flushed control group also underwent blastocyst recovery, handling and transfer, neither effect is attributable to superovulation, the transfer procedure or the recipient uterine environment.

This mirrors a consistent human pattern in which the mode of transfer rather than in vitro fertilisation itself shapes offspring size, and the strongest evidence comes from a design closely analogous to ours. In a Nordic registry study of 4,510,790 singletons that included 33,056 sibling groups conceived by at least two different methods, singletons born after fresh embryo transfer had lower mean birthweight and increased odds of being small for gestational age, whereas those born after frozen embryo transfer had higher mean birthweight and increased odds of being large for gestational age, in both cases relative to naturally conceived siblings^46^. The within-sibship comparison removes maternal and subfertility confounding in the same way that our in vivo-fertilised, transferred control removes the procedural contributions listed above, and both designs therefore isolate cryopreservation as the step associated with increased growth. The clinical significance lies less in birth size itself than in what a large birth predicts, since being born large for gestational age is associated with an increased risk of obesity in adulthood^47^. Taken with the observation that the growth effect here was sex-specific, this argues for attention to growth trajectory rather than birth weight alone.

### Reduced respiratory capacity with increased H_2_O_2_ production indicates a narrowed cardiac mitochondrial capacity

In the adult offspring left ventricle, vitrification reduced complex I- and convergent complex I+II-supported OXPHOS and maximal noncoupled electron transfer capacity, whereas IVF reduced LEAK respiration and fatty acid-supported respiration. Both procedures increased H_2_O_2_ production relative to oxygen flux across multiple respiratory states.

The heart derives the great majority of its ATP from OXPHOS and depends on metabolic flexibility to accommodate changes in workload and substrate supply^23^. A ventricle with reduced maximal capacity and elevated ROS production may be indistinguishable from control under basal conditions yet less resilient during ageing^48^, exercise^49^, pressure overload^50^ or ischaemia–reperfusion^28^. The reduction in fatty acid-supported OXPHOS in IVF offspring is relevant in the same way, because fatty acid oxidation becomes the dominant source of myocardial ATP after the perinatal metabolic transition^24^, so a lower intrinsic capacity to oxidise fatty acid-derived substrate constrains the pathway on which the adult heart principally relies.

This interpretation aligns with developmental programming models driven by entirely different insults. Prenatal hypoxia reduces aerobic capacity and raises ROS in adult murine cardiac mitochondria in a sex-dependent manner^27^, reduces respiratory chain subunit abundance and maximal oxygen consumption in guinea pig offspring^29^, and raises basal H_2_O_2_ production while increasing sensitivity to ischaemia–reperfusion^28^; a comparable signature is already present in the fetal heart^30^. Maternal undernutrition similarly lowers adult cardiac fatty acid oxidative capacity and maximal OXPHOS^26^. Embryo manipulation produces a comparable phenotype that places the periconceptional window alongside these later gestational insults as a route to cardiac mitochondrial programming, consistent with the broader principle that the periconceptional environment shapes lifelong health^3,18^.

Only one previous study has examined cardiac mitochondria after assisted conception, reporting reduced myocardial mitochondrial abundance and lower OXPHOS complex I and IV protein in male adolescent sheep following in vitro embryo culture^36^. The present data extend that observation from abundance to function, and show that the deficit is not confined to IVF but is produced independently by cryopreservation.

Whether the adult phenotype descends directly from the embryonic disturbance or arises secondarily from the cardiac remodelling cannot be resolved by these data. The same broad signature, depolarised mitochondria with elevated ROS, was present in the blastocyst and in the adult heart, which is suggestive of a single trajectory. However, concentric remodelling itself raises myocardial ATP demand, so the structural and mitochondrial phenotypes are unlikely to be independent. Distinguishing the two possibilities requires longitudinal measurement of cardiac mitochondrial function across the life course, since a defect that is present from the outset and stable would favour programming, whereas one that emerges in step with the geometric change would favour an acquired mechanism.

### Complex III is a plausible convergent site of functional vulnerability for both IVF and vitrification

In our study, complex III appeared to be a convergent site of functional vulnerability, being the only complex whose catalytic activity was reduced by both IVF and vitrification. The pharmacology supports this: between-group differences in H_2_O_2_ production persisted after complex I inhibition with rotenone, arguing against complex I and reverse electron transport as the principal source^25^, but were abolished by antimycin A, which blocks the Q_i_ site and drives all groups to a maximal Q_o_ site ROS-producing state^51^, implicating the Q_o_ site of complex III where ubiquinol oxidation yields a superoxide-generating ubisemiquinone^52^. These observations localise the defect by inference rather than by direct measurement, and targeted assessment of Q_o_ site ROS production would be required to establish it.

Complex III was also the point at which subunit abundance and catalytic activity diverged most clearly. UQCRC2 abundance increased while complex III activity fell, and the same relationship held for ATP5A and complex V. Altered respiratory chain assembly could explain this discordance, since complexes I, III and IV associate into supercomplexes whose assembly determines the route and efficiency of electron flux^53^. Subunits synthesised as a compensatory response that do not assemble into functional units would account for three otherwise disparate observations: raised abundance alongside reduced activity, increased ROS despite reduced respiration, and reduced complex II-supported electron transfer despite fully preserved complex II activity. The last is particularly informative, because complex II is not itself incorporated into the respirasome and its electrons must still pass through complex III^53^, so a downstream limitation at complex III would reduce complex II-supported respiration even when complex II is entirely intact.

Although both procedures converged on complex III, they arrived there from different directions. IVF reduced the activities of complexes III, IV and V and lowered fatty acid-supported respiration, whereas vitrification reduced the activities of complexes I and III and lowered coupled and maximal respiration. Across most endpoints there was no interaction between them, indicating that IVF and cryopreservation act through largely separate mechanisms rather than one modifying the effect of the other. This bears directly on the shift towards freeze-all strategies, which exposes an increasing proportion of ART-conceived children to both exposures: if the two steps act largely independently, neither can be assumed harmless on the grounds that the other is the dominant exposure, and each represents a distinct target for protocol optimisations.

## Limitations

Several limitations should be acknowledged. IVF-derived embryos were necessarily cultured in vitro before assessment or transfer, so the IVF arm reflects the combined contribution of the fertilisation method and the culture environment, which cannot be separated within this design. Offspring within a litter are not statistically independent, and because IVF itself reduced litter size, litter represents both a clustering variable and a potential mediator of the growth phenotype, yet offspring were treated as independent observations. Sex was likewise not included as a factor in the cardiac and mitochondrial analyses, although the growth effects reported here were sex-specific and developmental programming of cardiac mitochondria is frequently sex-dependent^27,29,30^, so effects confined to one sex may have been diluted. Mitochondrial function was assessed in isolated organelles, which gives direct access to intrinsic respiratory properties but removes cytosolic substrate supply, calcium handling, network organisation and the workload imposed in vivo; immunoblotting assessed a single representative subunit of each complex and therefore cannot report assembly intermediates or respiratory supercomplex organisation, which matters because altered assembly was invoked to explain the dissociation between subunit abundance and catalytic activity, and, for the same reason, localisation of the excess H_2_O_2_ to complex III rests on pharmacological inference rather than direct measurement at the Q_o_ site. Finally, offspring were studied at a single adult age and under unstressed conditions. This is the most important constraint on the central interpretation, because a narrowed metabolic reserve is by definition not apparent at rest: establishing whether it carries a functional cost requires challenge by exercise, ageing, pressure overload or ischaemia–reperfusion, and establishing when it arises requires measurement across the life course.

## Conclusion and future perspectives

Our study demonstrates that IVF and vitrification reduced respiratory capacity and increased H_2_O_2_ production in adult offspring, with complex III as a convergent site of functional vulnerability for both procedures. The same disturbance in mitochondrial and redox homeostasis was present in developmentally competent blastocysts that went on to establish pregnancies. These findings indicate that in vitro culture and cryopreservation contribute through largely independent routes.

Whether the adult phenotype is programmed from the embryo or acquired alongside the cardiac remodelling remains the central open question, and longitudinal measurement across the life course would resolve it. The role of complex III should be tested directly, through targeted assessment of Q_o_ site ROS production and of respiratory supercomplex assembly^53^. Most importantly, a narrowed metabolic reserve is by definition invisible at rest, so establishing whether this phenotype carries a physiological cost requires challenging these hearts by exercise, ageing, pressure overload or ischaemia–reperfusion, in a sex-stratified design^27,29,30^. If early oxidative stress is the initiating event, antioxidant supplementation of culture and vitrification media is an obvious target for protocol refinement^34^.

Collectively, our results show that IVF and vitrification can disrupt preimplantation embryo redox balance and remodel the cardiac structure and metabolism of adult mouse offspring. Mitochondrial and redox function was disturbed at both stages, and may therefore represent one route by which the periconceptional environment comes to influence long-term cardiac health.

## Acknowledgements

This study was funded by a BBSRC CASE PhD studentship

